# CoMut: Visualizing integrated molecular information with comutation plots

**DOI:** 10.1101/2020.03.18.997361

**Authors:** Jett Crowdis, Meng Xiao He, Brendan Reardon, Eliezer M. Van Allen

## Abstract

**Motivation:** Large-scale sequencing studies have created a need to succinctly visualize genomic characteristics of patient cohorts linked to widely variable phenotypic information. This is often done by visualizing the co-occurrence of variants with comutation plots. Current tools lack the ability to create highly customizable and publication quality comutation plots from arbitrary user data.

**Results:** We developed CoMut, a stand-alone, object-oriented Python package that creates comutation plots from arbitrary input data, including categorical data, continuous data, bar graphs, side bar graphs, and data that describes relationships between samples.

**Availability and Implementation:** The CoMut package is open source and is available at https://github.com/vanallenlab/comut under the MIT License, along with documentation and examples. A no installation, easy-to-use implementation is available on Google Colab (see GitHub).

## 1 Introduction

A common representation of cohort-level mutation data in large-scale sequencing studies is a comutation plot, which shows sample level mutation status along with other relevant clinical and genomic characteristics. Originally designed in 2011 (Stransky *et al.*, 2011), comutation plots are now ubiquitous and provide a way to communicate mutations and other patterns in a cohort. Software currently exists to create comutation plots: CoMutPlotter (Huang *et al.*, 2019) and jsComut (Pearce *et al.*, 2019) provide a web interface for creating comutation plots while larger bioinformatic packages often have a function for creating comutation plots, including *oncoplot* in maftools (Mayakonda *et al.*, 2018) and *waterfall* in GenVisR (Skidmore *et al.*, 2016). However, these software only plot specific genomic and phenotypic data types, and as clinically integrated sequencing uses rapidly rise, comutation software that can capture the complexity of advances in molecular analyses and phenotypic information is necessary.

## 2 Methods

Here we present a Python package named ‘CoMut’ to streamline the creation of comutation plots. Implemented in Python’s plotting library, matplotlib, it is the first of its kind to utilize an object-oriented framework for maximum customizability. As a result, users can freely edit individual parts of the plot after its creation and fine-tune plots for publication. Furthermore, instead of being constrained to a limited number of prespecified genomic data types, CoMut is data agnostic and enables the plotting of arbitrary data types. For example, categorical data encompasses mutation data (e.g. variant types) and most clinical variables (e.g. tumour stage). Categorical data are drawn as boxes with user specified colors, and two mutations in the same gene within one sample are drawn as triangles rather than boxes. This allows CoMut to depict allele-specific copy number alterations or plot mutations and copy number alterations together, which are both major advantages relative to existing software. CoMut also supports continuous data, bar graphs (e.g. mutation burden), side bar graphs, and sample indicators, which indicate relationships between samples.

CoMut uses the Pandas library to handle data and accepts a variety of file types, including tsv, csv, and maf file formats. CoMut includes helper functions to parse common file types into dataframes, and it can export plots in both raster (.jpg, .png) and vector (.pdf, .svg) forms. It can also handle missing data, an important feature for clinical sequencing studies where some data types for individuals may be unavailable. We provide a quickstart notebook in GitHub connected to Google Colab that allows users to create basic comutation plots from maf files using their browser without any installations.

## 3 Usage Scenario

To illustrate the features of CoMut, we created a comutation plot visualizing a cohort from a study of selective response to immunotherapy in melanoma (Liu *et al.*, 2019) (Figure 1). We obtained mutation and clinical data from the supplement and used allele-specific copy number profiles from ABSOLUTE (Carter *et al.*, 2012) to classify allele-specific copy number alterations. In brief, we defined samples as whole genome doubled if they had an average ploidy greater than 2.5. We classified copy number alterations in genes by comparing the integer copy number of the segment on which the gene was located to baseline allelic ploidy (2 if a sample was WGD, 1 otherwise).

**Fig. 1.**
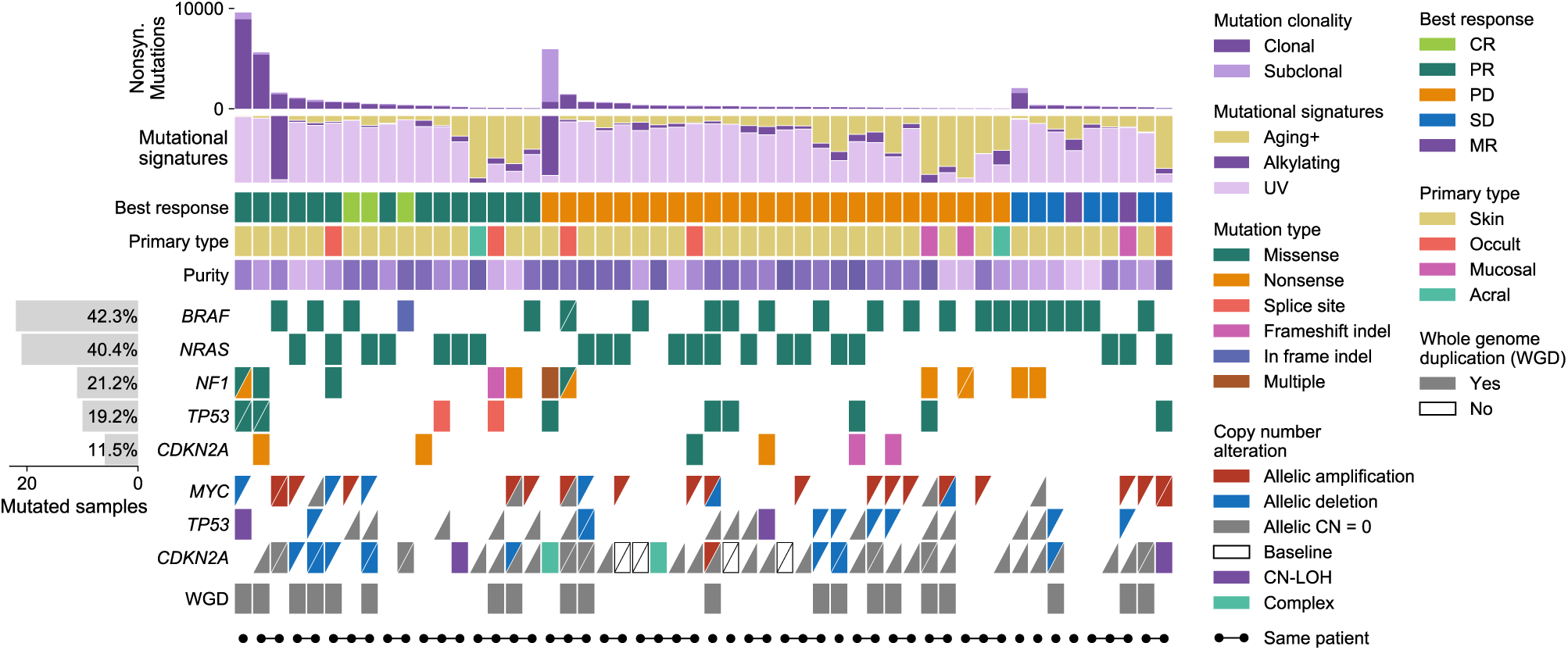
A comutation plot generated with CoMut using data provided in Liu et al., 2019. For visualization purposes, only 52 samples are shown. Each column represents a tumour. Tumours are ordered by best RECIST criteria response (CR, PR, PD, SD, or MR) and within each subgroup by nonsynonymous mutation load. Allele-specific copy number data are shown as triangles and classified relative to reference ploidy (2 if sample has whole genome duplication, 1 otherwise). Unfilled boxes with a slash indicate that data was unavailable due to low coverage. *Complex* indicates that the gene fell into two segments of different copy number. CN-LOH indicates copy neutral loss of heterozygosity. Sample indicators are added for demonstration purposes and do not represent data from the study.

To create the comutation plot, we added the following datasets to the plot using CoMut, specifying color mappings for each. Mutation type, copy number alterations, and discrete clinical data (e.g. primary type) were all added as categorical data. Purity was added as continuous data, and mutation burden and mutational signatures were added as bar graphs. We calculated the number of samples mutated in each gene and added this data as a side bar graph. We added sample indicators for demonstration, though they do not represent data from the study. All of this was completed using CoMut’s built-in functions for adding specific data types and was achieved with only a few lines of code. Leveraging CoMut’s object-oriented framework, small edits to the figure were easily made by acting on specific subplots after the plot was made (for example, moving the y-axis labels inside the side bar graph). The legend was constructed using CoMut’s legend functions. This visualization reveals that allelic copy number alterations are common and that *CDKN2A* and *TP53* often experience deletions after whole genome duplication in this study. The data and code to create this comutation plot can be found in the CoMut GitHub repository.

## 4 Conclusion

CoMut is a highly customizable tool for creating comutation plots to visualize arbitrary genomic and clinical characteristics of samples in sequencing studies. It supports a variety of data types and allows the user complete control over the structure and appearance of the plot. Its object-oriented framework allows users to customize the plot for publication and allows developers to extend CoMut’s functionality. By providing a quick-start notebook integrated with Google Colab, we also provide an easy way for those without programming experience to create comutation plots using only input files and a browser.

## Acknowledgements

We acknowledge and thank Van Allen lab members Daniel Boiarsky, Seunghun Han, and Cora Ricker for testing the package and providing feedback.

